## Supporting information (Fig S1-S2, Tables S3-S6) for "Genome-wide analysis of fitness determinants of *Staphylococcus aureus* during growth in milk"

### Supporting figures

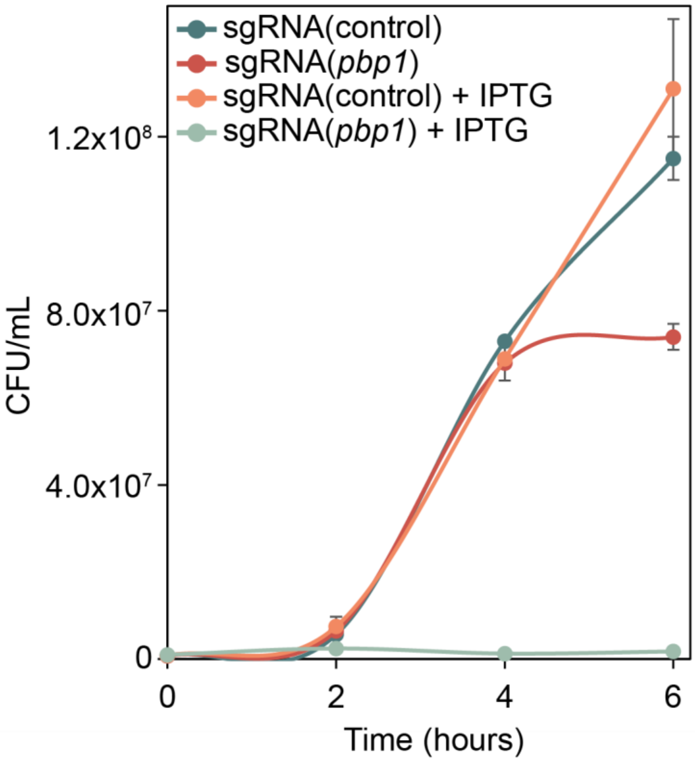


**Fig. S1.** CFU/ml of CRISPRi strains targeting *pbp1* (MK1857) and a control strain harboring a non-targeting sgRNA (MM75). Strains were grown in UHT milk for 6 hours with or without induction with 500 µm IPTG. CFU/ml was calculated at 2-hour intervals. The data represent the average of two independent experiments, with error bars indicating standard error.

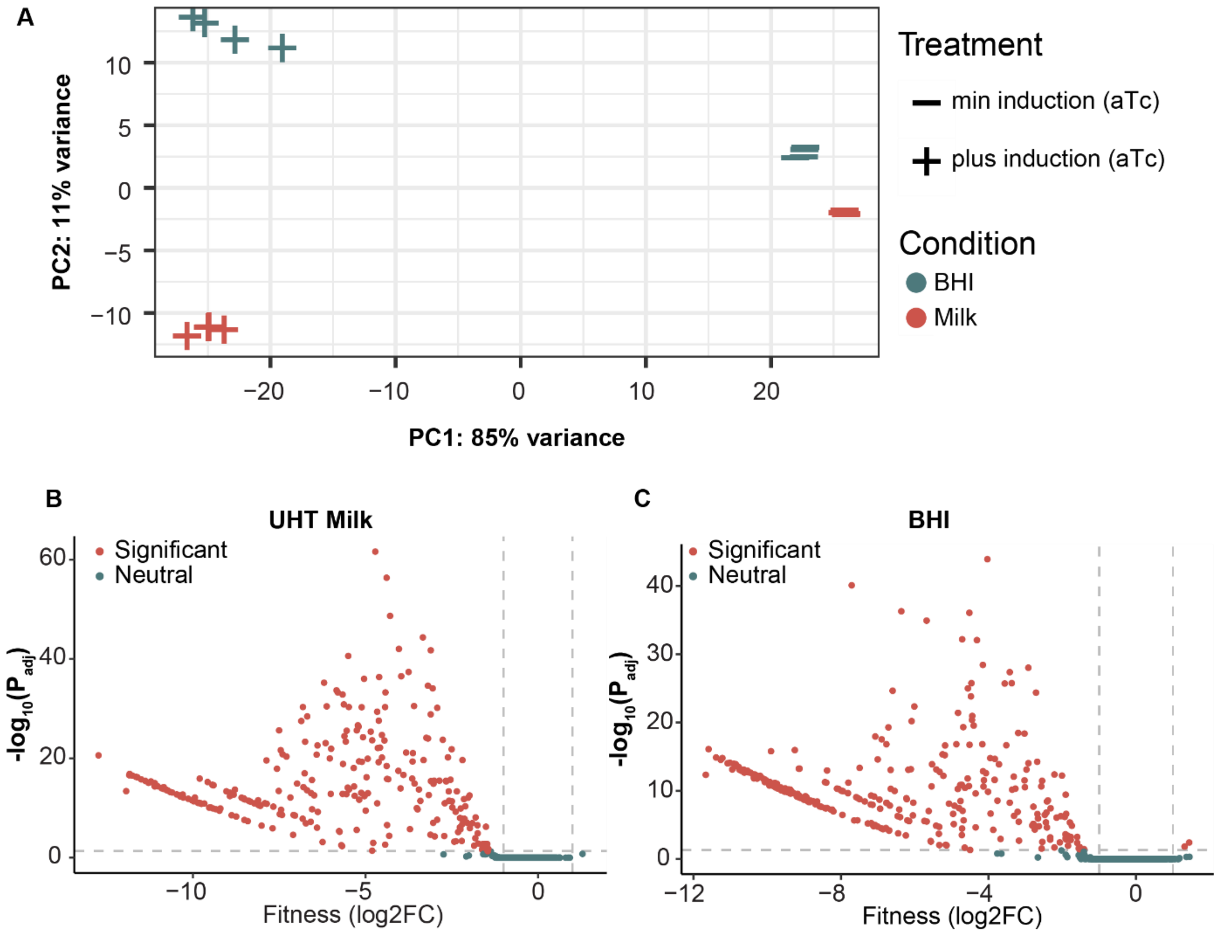


**Fig. S2**. **(A)** Principal component analysis (PCA) of the rlog-transformed sgRNA counts. **(B)** Fitness effect upon dCas9 induction in UHT milk. **(C)** Fitness effect upon dCas9 induction in BHI. Genes with significant fitness effects (|log2FC| ≥ 1, *Padj* < 0.05) are depicted in red, while green points indicate neutral genes. Dashed grey lines denote the thresholds for significance (|log2FC| ≥ 1, *Padj* < 0.05). The data were analyzed using DESeq2 [1], log2FC values were shrunk using the 'apeglm' method [2]

### Supporting tables

**Table S1. CRISPRi screen to identify genes influencing fitness in milk versus BHI.** Log2 fold change (L2FC) and the adjusted p-value (padj) and the conclusion about significance (“essential” or “costly” if |L2FC| ≥ 1, $P_{adj}$ 0.05) is shown for each sgRNA. Interaction effect is also indicated, representing the differential effect of aTc induction in milk vs. BHI, and whether this is significant (“yes” if |L2FC| ≥ 1, $P_{adj} < 0.05$).

**Table S2. CRISPRi screen to identify genes influencing TMP-SMX susceptibility**. Log2 fold change (L2FC) and the adjusted p-value (padj) and the conclusion about significance (“essential” or “costly” if |L2FC| ≥ 1, $P_{adj}$ 0.05) is shown for each sgRNA in treated and untreated samples. Interaction effect is also indicated, representing the differential effect of aTc induction in the presence of TMP-SMX compared to untreated samples, and whether this is significant (“yes” if |L2FC| ≥ 1, $P_{adj} < 0.05$).

**Table S3.** List of genes less important for fitness in milk as determined by CRISPRi-seq

| **Locus tag targeted (SAOUHSC)** | **Target gene(s)**[a] | **Function/pathway** [b] | **Interaction $log_2FC$** [c] | **Interaction $P_{adj}$** [c] |
|---|---|---|---|---|
| _00867 | *dltXABCD* | D-alanylation of teichoic acids | 8.7 | 2,81E-08 |
| _02399 | *glmS* | Glucosamine-fructose-6-phosphate aminotransferase | 7.7 | 2,4E-06 |
| _02368 | *pyrG* | CTP synthase | 7.5 | 5,29E-06 |
| _00561 | *vraX* | Cell wall stress protein | 6.9 | 1,04E-30 |
| _00889 | *mnhABCDEFG* | Monovalent cation/H+ antiporter | 6.9 | 4,29E-05 |
| _01285 | *glnRA* | Glutamine synthetase repressor-Glutamine synthase | 6.0 | 0,000833 |
| _00781 | *hprK*-*lgt*-00783-00784-*trxB* | Lipoprotein | 6.0 | 2,08E-05 |
| _00953 | *ugtP*-*ltaA* | Glycolipid/Lipoteichic acid biosynthesis | 6.0 | 1,36E-11 |
| _01895 | *sagB* | Putative β-N-acetylglucosaminidase | 5.8 | 1,36E-19 |
| _00906 | 00906 | Fumarylacetoacetate hydrolase | 5.7 | 9,57E-05 |
| _01702 | *mtnN*-01701-01700-*aroE*-01698-*nad*-01696-*rsfS* | Chorismate biosynthesis | 5.7 | 0,002688 |
| _00934 | *spxA* | Transcriptional regulator | 5.4 | 0,000245 |
| _00788 | 00788 | GlmS regulation | 5.0 | 0,000436 |
| _00920 | *fabHF* | | 4.7 | 0,036216 |
| _01106 | *murI*-01107-01108 | Glutamate racemace | 4.5 | 0,000521 |
| _02801 | *gtaB* | UTP-glucose-1-phosphate uridylyltransferase | 4.3 | 7,83E-08 |
| _02793 | *pgcA* | Phosphoglucosamine mutase | 4.2 | 2,39E-12 |
| _02997 | 02997-*msrA3* | Acetyltransferase -ethionine-sulfoxide reductase | 4.2 | 4,88E-18 |
| _00762 | *tagO* | Wall teichoic acid biosynthesis | 4.1 | 0,012117 |
| _01622 | 1622-*nusB*-*xseA*-1619-*ispA* | Endoribonuclease | 4.0 | 3,64E-07 |
| _00980 | *menA* | Menaquinone biosynthesis | 3.9 | 2,4E-06 |
| _02369 | *rpoE* | RNA polymerase, subunit | 3.8 | 0,000537 |
| _01265 | 01265 | Metallophosphoesterase | 3.7 | 2,19E-05 |
| _00567 | 00567 | Membrane protein | 3.6 | 0,040001 |
| _02612 | *rpiA* | Ribose-5-phosphate isomerase A | 3.4 | 1,25E-09 |
| _00189 | 0189-0190 | Membrane protein | 3.4 | 4,2E-09 |
| _A02795 | A02795 | | 3.4 | 7,11E-06 |
| _02121 | 02121 | CamS pheromone | 3.3 | 3,77E-20 |
| _01746 | *secDF* | Preprotein translocase | 3.2 | 1,18E-08 |
| _02366 | *fbaA* | Fructose-bisphosphate aldolase | 3.1 | 0,000109 |
| _00640 | *tagA* | Wall teichoic acid biosynthesis | 3.1 | 0,004214 |
| _00474 | *rplY* | | 3.0 | 1,91E-05 |
| _01154 | *sepF* | Cell division factor | 3.0 | 4,15E-16 |
| _01482 | *aroB* | Chorismate biosynthesis | 3.0 | 1,08E-05 |
| _01481 | *aroA* | Chorismate biosynthesis | 3.0 | 7,47E-05 |
| _00832 | *aroD* | Chorismate biosynthesis | 3.0 | 1,43E-09 |
| _01809 | *accDA* | Fatty acid biosynthesis | 2.9 | 0,000108 |

| | | | | |
|---|---|---|---|---|
| _01495 | 01495 | | 2.9 | 3,7E-13 |
| _01586 | *srrAB* | Two-component system | 2.9 | 8E-05 |
| _00964 | *0964-0965* | CPBP family intra-membrane metalloprotease | 2.8 | 1,42E-08 |
| _01223 | *gid* | tRNA methyltransferase | 2.6 | 0,000615 |
| _02611 | *lyrA* | CPBP family intra-membrane metalloprotease | 2.6 | 7,47E-05 |
| _01852 | *aroA2* | Chorismate biosynthesis | 2.4 | 1,39E-06 |
| _00253 | 00253 | | 2.3 | 0,021076 |
| _01635 | *aroK* | Chorismate biosynthesis | 2.3 | 2,91E-05 |
| _00996 | 00996 | | 2.1 | 0,010107 |
| _00787 | *rapZ-00788* | GlmS regulation | 2.1 | 7,55E-08 |
| _01022 | *thiV* | Thiamine ABC transporter | 2.0 | 1,96E-05 |
| _01405 | 01405 | | 1.9 | 0,00587 |
| _02143 | 02143 | | 1.9 | 0,036216 |
| _01501 | *ebpS* | Elastin binding protein | 1.8 | 8,56E-05 |
| _01821 | 01821 | DNA methylase | 1.8 | 0,016802 |
| _02899 | *02899-02900* | Aminohydrolase | 1.6 | 0,025009 |
| _01480 | 01480-01479-01478 | | 1.6 | 0,03361 |

[a] Genes in the same operon as the target gene is also indicated.

[b] Functional characterization from *Aureo*Wiki [3].

[c] L2FC ($log_2$fold change) in fitness upon CRISPRi depletion and adjusted *p*-values ($P_{adj}$) from DESeq2 analysis

**Table S4.** Strains used in this study

| Strain | Genotype and Characteristics | Reference |
|---|---|---|
| ***E. coli*** | | |
| IM08B | DH10B, Δ*dcm*, $P_{help}$-*hsdMS*, $P_{N25}$-*hsdS* (strain expressing the *S. aureus* CC8 specific methylation genes) | [4] |
| ***S. aureus*** | | |
| NCTC8325-4 | Derivative of NCTC8325, cured of prophages | [5] |
| MH225 | NCTC8325-4, pLOW-Pspac2-*dcas9*, $ery^r$ | [6] |
| MK1857 | MH225, pCG248-sgRNA(*pbp1*), $ery^r$, $cam^r$ | This work |
| MM75 | MH225, pCG248-sgRNA(*luc*). $ery^r$, $cam^r$ | [6] |
| MM223 | NCTC8325-4, *tetR*-Ptet-*dcas9* | This work |
| MM230 | MM223, pCG248-sgRNA(*luc*), $cam^r$ | This work |
| MM267 | MM223, *tetM* | This work |
| MM268 | MM267, pCG248sgRNA(*luc*), $cam^r$ | This work |
| MM269 | MM267, pCG248-sgRNA(*pbp1*), $cam^r$ | This work |
| MM289 | MM267, pVL2336-sgRNA(*sarA*), $cam^r$ | This work |
| MM290 | MM267, pVL2336-sgRNA(*nrdF*), $cam^r$ | This work |
| MM294 | MM267, pVL2336-sgRNA(*purE*), $cam^r$ | This work |
| MM295 | MM267, pVL2336-sgRNA(*purA*), $cam^r$ | This work |
| MM312 | MM267, pVL2336-sgRNA(*purB*), $cam^r$ | This work |
| MM313 | MM267, pVL2336-sgRNA(*thyA*), $cam^r$ | This work |
| MM314 | MM267, pVL2336-sgRNA(*fhuC*), $cam^r$ | This work |
| MM315 | MM267, pVL2336-sgRNA(*htsA*), $cam^r$ | This work |
| MM316 | MM267, pVL2336-sgRNA(*mntA*), $cam^r$ | This work |
| MM318 | MM267, pVL2336-sgRNA(*sucC*), $cam^r$ | This work |
| MM342 | MM267, pVL2336-sgRNA(*clpP*), $cam^r$ | This work |
| MM403 | MM267, pVL2336-sgRNA(*noc*), $cam^r$ | This work |
| MM404 | MM267, pCG248-sgRNA(*SAOUHSC_01782*), $cam^r$ | This work |
| MM421 | MM267, pVL2336-sgRNA(polA), $cam^r$ | This work |
| MM422 | MM267, pVL2336-sgRNA(*nupC*), $cam^r$ | This work |
| MM423 | MM267, pVL2336-sgRNA(*nupG*), $cam^r$ | This work |
| MM424 | MM267, pVL2336-sgRNA(*murB*), $cam^r$ | This work |
| MM425 | MM267, pVL2336-sgRNA(*ung*), $cam^r$ | This work |
| MM426 | MM267, pVL2336-sgRNA(*SAOUHSC_02121*), $cam^r$ | This work |

$cam^r$: chloramphenicol resistance; $ery^r$: erythromycin resistance

**Table S5.** Plasmids used in this study

| Plasmid | Description | Reference |
|---|---|---|
| pCG248 | *E. coli/S. aureus* shuttle vector, amp$^r$, cam$^r$ | [7] |
| pCG248-sgRNA(non-target) | For constitutive expression of sgRNA, non-targeting, amp$^r$, cam$^r$ | [7] |
| pCG248-sgRNA(SAOUHSC_01782) | For constitutive expression of sgRNA, amp$^r$, cam$^r$ | This work |
| pCG248-sgRNA(*pbp1*) | For constitutive expression of sgRNA, amp$^r$, cam$^r$ | [7] |
| pVL2336 | *E. coli/S. aureus* shuttle vector, amp$^r$, cam$^r$ | [6] |
| pVL2336-sgRNA(*nrdF*) | For constitutive expression of sgRNA, amp$^r$, cam$^r$ | [6] |
| pVL2336-sgRNA(*noc*) | For constitutive expression of sgRNA, amp$^r$, cam$^r$ | This work |
| pVL2336-sgRNA(*purA*) | For constitutive expression of sgRNA, amp$^r$, cam$^r$ | This work |
| pVL2336-sgRNA(*purE*) | For constitutive expression of sgRNA, amp$^r$, cam$^r$ | This work |
| pVL2336-sgRNA(*clpP*) | For constitutive expression of sgRNA, amp$^r$, cam$^r$ | This work |
| pVL2336-sgRNA(*sarA*) | For constitutive expression of sgRNA, amp$^r$, cam$^r$ | This work |
| pVL2336-sgRNA(*purB*) | For constitutive expression of sgRNA, amp$^r$, cam$^r$ | This work |
| pVL2336-sgRNA(*thyA*) | For constitutive expression of sgRNA, amp$^r$, cam$^r$ | This work |
| pVL2336-sgRNA(*fhuC*) | For constitutive expression of sgRNA, amp$^r$, cam$^r$ | This work |
| pVL2336-sgRNA(*htsA*) | For constitutive expression of sgRNA, amp$^r$, cam$^r$ | This work |
| pVL2336-sgRNA(*mntA*) | For constitutive expression of sgRNA, amp$^r$, cam$^r$ | This work |
| pVL2336-sgRNA(*sucC)* | For constitutive expression of sgRNA, amp$^r$, cam$^r$ | This work |
| pVL2336-sgRNA(*polA*) | For constitutive expression of sgRNA, amp$^r$, cam$^r$ | This work |
| pVL2336-sgRNA(*nupC*) | For constitutive expression of sgRNA, amp$^r$, cam$^r$ | This work |
| pVL2336-sgRNA(*nupG*) | For constitutive expression of sgRNA, amp$^r$, cam$^r$ | This work |
| pVL2336-sgRNA(*murB*) | For constitutive expression of sgRNA, amp$^r$, cam$^r$ | This work |
| pVL2336-sgRNA(*ung*) | For constitutive expression of sgRNA, amp$^r$, cam$^r$ | This work |
| pVL2336-sgRNA(SAOUHSC_02121) | For constitutive expression of sgRNA, amp$^r$, cam$^r$ | This work |
| pMAD | Vector for allelic replacement, amp$^r$, ery$^r$ | [8] |
| pMAD-GG | pMAD adapted for Golden Gate cloning, amp$^r$, ery$^r$ | [6] |
| pFD152 | *E. coli/S. aureus* shuttle vector, *tetR*-Ptet-*dcas9* | [9] |
| pCN36 | *E. coli/S. aureus* shuttle vector, *tetM* | [10] |

cam$^r$: chloramphenicol resistance; ery$^r$: erythromycin resistance; amp$^r$: ampicillin resistance

**Table S6.** Oligos used in this study

| Oligo name | Sequence (5' – 3') |
|---|---|
| **Construction of pMAD-*tetR*-Ptet-*dcas9*** | |
| mm40_GG_ori_up_F | GTGTCTGGTCTCCCTATCCAAGCAGTTAACGTACAAAC |
| mm32_GG_ori_up_R | GTCCAAGGTCTCGCATCGAACCCCGATGTTGTC |
| mm33_GG_Ptet_dCas9_tetR_F | GTCCAAGGTCTCCGATGCTTTTAAGACCCACTTTCAC |
| mm34_GG_Ptet_dCas9_tetR_R | GTCCAAGGTCTCCGTCATAAACGCAGAAAGGCCCAC |
| mm35_GG_ori_down_F | GTCCAAGGTCTCCTGACTCCCTTAAGACAGACCTG |
| mm36_GG_ori_down_R | GTCCAAGGTCTCCCTGCCATTGGTGGTATCGCTGTTG |
| **Construction of individual sgRNAs** | |
| *purA*_F | TATACTCCGCCAAATTGAATGGTA |
| *purA*_R | AAACTACCATTCAATTTGGCGGAG |
| *purB*_F | TATATAGTAAAGGCTACAACATCA |
| *purB*_R | AAACTGATGTTGTAGCCTTTACTA |
| *purE*_F | TATAGATACTACTTGTTTTTCGTA |
| *purE*_R | AAACTACGAAAAACAAGTAGTATC |
| *thyA*_F | TATAACTTTCTTTGTCGTTAATAG |
| *thyA*_R | AAACCTATTAACGACAAAGAAAGT |
| *sarA*_F | TATATTAACTGCTTTAACAACTTG |
| *sarA*_R | AAACCAAGTTGTTAAAGCAGTTAA |
| *htsA*_F | TATAACAAGCTGCAACTAAAAGTA |
| *htsA*_R | AAACTACTTTTAGTTGCAGCTTGT |
| *fhuC*_F | TATAATTGACGTCACTTTGCCATC |
| *fhuC*_R | AAACGATGGCAAAGTGACGTCAAT |
| *hemQ*_F | TATAATACCAACCATCTAATGTTT |
| *hemQ*_R | AAACAAACATTAGATGGTTGGTAT |
| *mntA*_F | TATAGCCGCGTACTGGTATCGATA |
| *mntA*_R | AAACTATCGATACCAGTACGCGGC |
| *nupC*_F | TATAAAGAAGAATGGTGGTTGCTT |
| *nupC*_R | AAACAAGCAACCACCATTCTTCTT |
| *nupG*_F | TATATAAAGAACCATGCTAAAAAC |
| *nupG*_R | AAACGTTTTTAGCATGGTTCTTTA |
| *polA*_F | TATATGCAAAACCATATACTGCAT |
| *polA*_R | AAACATGCAGTATATGGTTTTGCA |
| *noc*_F | TATAAACGATACGTTCAATTTGAA |
| *noc*_R | AAACTTCAAATTGAACGTATCGTT |
| *ung*_F | TATAGATATATATTTTCCCTATCA |
| *ung*_R | AAACTGATAGGGAAAATATATATC |
| *murB*_F | TATAGTATAAGTGTATCGTTTTAA |
| *murB*_R | AAACTTAAAACGATACACTTATAC |
| SAOUHSC_01782_F | TATATCAAAAATATTTTGACCTTC |
| SAOUHSC_01782_R | AAACGAAGGTCAAAATATTTTTGA |
| SAOUHSC_02121_F | TATATTCCAGCCTGGTCATCCTTA |
| SAOUHSC_02121_R | AAACTAAGGATGACCAGGCTGGAA |
